# EnVhogDB: an extended view of the viral protein families on Earth through a vast collection of HMM profiles

**DOI:** 10.1101/2024.06.25.600602

**Authors:** Pérez-Bucio Rubén, François Enault, Clovis Galiez

**Author notes:** these authors equally contributed to the work (François Enault, Clovis Galiez).

## Abstract

Over the last twenty years, hundreds of metagenomic studies have generated millions of viral genomic sequences from a wide variety of ecosystems. Despite this, the overall genetic diversity of viruses remains elusive, both in terms of the number of protein families they encode and the diversity of these families. Indeed, even if it is recognized that the organization of the viral protein sequence space requires sensitive homology detection methods, such methods have never been applied at a large scale. To produce a more realistic and comprehensive view of the protein diversity in the viral world, we have (i) collected thousands of viromes and identified viral contigs and proteins within them, (ii) retrieved viral proteins available in different public databases, and (iii) applied sensitive similarity searches to cluster all these proteins into families and (iv) annotated the protein clusters produced. More than 46 million deduplicated proteins were clustered into less than 2.3 million protein families. After further removing genomic sequences likely of cellular origin using an iterative procedure, the remaining 2,203,457 clusters were coined enVhogs (for environmental Viral homologous groups). Their multiple sequence alignments have been transformed into HMMs to constitute the EnVhog database. Even if only a small proportion of enVhogs were annotated (15.9 %), they encompass almost half of the protein dataset (44.8 %). Applied to the annotation of four recently published viromes from diverse environments (sulfuric soil, grassland, surface seawater and human gut), enVhog HMMs doubled the number of viral sequences characterized, and increased by 54%-74% the number of proteins functionally annotated. EnVhogDB, the largest comprehensive compilation of viral protein information to date, is a resource that will thus further help to determine the functions of proteins encoded in newly sequenced viral genomes, and help to improve the accuracy of viral sequence detection tools.

EnVhog database is available at http://envhog.u-ga.fr/envhog.

## Introduction

With the remarkable estimated amount of at least 10^31^ viral particles on the planet, viruses are probably the most abundant and diverse biological entity on Earth (Mushegian 2020). Viruses can be found in almost all studied ecosystems, ranging from ocean samples to host-associated microenvironments such as the human gut (Cobián Güemes et al. 2016). Viruses are key players in all these ecosystems, from driving cellular mortality and evolution, to influencing global-scale biogeochemical cycles (Wieczynski et al. 2023). In addition to their large number, viruses are diverse, as evidenced by the existence of numerous groups with independent origins (Koonin et al. 2022). The many strategies implemented by these entities are reflected in the nature of their genetic support and also in the size of their genomes, which ranges from 1.7 Kb to 2.3 Mb (Opriessnig et al. 2020; Philippe et al. 2013). Thus, despite their small individual size, the number of different protein families encountered in viruses is high, especially as many cellular genes, even ribosomal ones, have been sporadically incorporated into viral genomes (Mizuno et al. 2019). Moreover, each viral protein family is diverse with high sequence heterogeneity (Terzian et al. 2021).

Due to this diversity in terms of number of genome families, protein families and of the heterogeneity of most of these families, viral sequence space still remains far from well characterized. Virus genomes are accessed through isolation on a host for more than 40 years. Yet, as only a small fraction of microbes have culture representatives, most viruses on Earth remain uncultivated. To discover previously unseen genomes of viruses, metagenomic sequencing is the most commonly used methodology (Rosario and Breitbart, 2011). High throughput sequencing can be applied to study (i) microbiomes, by considering all microbes including viruses, or (ii) viromes by specifically targeting viruses mostly through prior filtration. Whether from microbial or viral metagenomes, the viral origin of the contigs must be verified as even viromes often contain a significant amount of cellular DNA (Roux et al. 2013). The identification of viral contigs from these datasets remains a difficult task as (i) some proteins are shared by the viral and cellular world and (ii) many proteins on contigs of viral origin are not similar to any known viral sequence (Roux et al. 2015a). Considering this last point, no homologs are detected for many new viral proteins because these proteins are too distantly related from known ones or because they correspond to unseen proteins. Without identified homologs, no functional annotation is possible for such viral proteins.

To improve this crucial goal of identifying homologs for new viral proteins, two avenues can be explored. First, the most straightforward solution is to use the largest possible database of viral proteins. Several viral sequence databases have been developed in recent years, made of either cultivated viruses (RefSeqVirus) or non-cultivated viral genomes from metagenomes of marine systems with efam (Zayed et al. 2021), or from all ecosystems such as GL-UVAB (Coutinho et al. 2019) or IMG/VR (Camargo et al. 2023). The second solution to better identify similarities between sequences of interest and reference proteins is to use remote homology detection methods. The most sensitive method that can be used routinely is based on Hidden Markov Model (HMM) comparisons (Söding 2005). Many viral protein HMM databases exists, here again either based on viral isolates such as POGs/pVOGs (Kristensen et al 2013; Grazziotin et al 2017), VOGDB (http://vogdb.org/) and PHROG (Terzian et al. 2021) or (ii) based on viral reference sequences and curated viral contigs from metagenomes, such as VPF (Paez-Espino et al, 2016) or efam (Zayed et al. 2021). Although these databases provide protein families in the form of HMMs, the creation of these families did not involve any remote homology detection step, with the exception of PHROG. To build these PHROGs, viral proteins were first clustered based on BLAST-like similarity searches as the other protein family databases. Then, HMMs of these protein clusters were generated and compared to each other using HMM-HMM comparisons and further used to build super-clusters (Terzian et al. 2021). This two step strategy proved to find more distant homologous relationships and result in generating more diverse protein families (Olo Ndela et al. 2021; Olo Ndela et al. 2022).

Here, we developed EnVhogDB, a database of viral protein families, whose interest lies both (i) in the incorporation of viral protein sequences from any possible sources, with an effort to collect all sequences from published viromes and (ii) in the clustering of these sequences using remote homology in the form of profile-sequence clustering. Note that the total database will be hereafter termed EnVhogDB, while enVhog will refer to a single cluster of homologous proteins.

## Materials & Methods

### Collecting viral contigs from in-house assembled public viromes

Scientific articles published between 2006 and 2019 dealing with viral metagenomics were identified using keyword searches in pubMed and google Scholar, and virome accession numbers were manually extracted from articles when possible. A total of 3,257 viromes were collected from these publications and downloaded from various databases, all as raw reads except for two that were already assembled. Each virome was downloaded, reads were trimmed and assembled, using softwares adapted to the sequencing technologies used (Table S1). In order to detect viral sequences among these viromes, contigs longer than 1.5 Kb were processed by VirSorter2 (Guo et al. 2021), VIBRANT (contigs containing at least four ORFs) (Kieft et al. 2020) and DeepVirFinder (score > 0.9 and p-value < 0.05) (Ren et al. 2020). Considering contigs predicted as viral by at least two of these three softwares, proteins were predicted using prodigal (v2.6.3; Hyatt et al. 2010) and only full-length proteins with less than five unknown amino acids (Xs) were considered here.

### Collecting viral genomic information and proteins from existing databases

In addition, viral genomic information was collected from multiple existing viral sequence databases. Reference genomes of (pro-)viruses were downloaded from PHROG (Terzian *et al*. 2021) and RefSeqVirus (accessed at 17/10/2022). Viral genomic sequences (*i.e*. complete genomes and genome fragments) obtained through culture-independent approaches, such as metagenomics, fosmid libraries, single-virus sequencing and prophage mining were also downloaded from different databases. Among these, IMG/VR (v4.1, Camargo et al. 2023) is mostly made of viral sequences extracted from microbiomes, GL-UVAB (Coutinho *et al*. 2019) mostly from marine viromes and microbiomes and efam (Zayed *et al*. 2021) only from marine viromes. Proteins encoded by these genomes and contigs were retrieved and added to virome proteins. All contigs were affiliated to a taxonomy using the MMseqs2 taxonomy tool, based on the taxonomic affiliation of RefSeqVirus sequences and using default parameters (Mirdita et al. 2021).

### Clustering all proteins

As some genomic sequences may be present in several databases, proteins were first deduplicated using the *linclust* module of MMseqs2 modules (Steinegger and Soeding 2018; Mirdita et al. 2019) with a minimum identity of 99% and a coverage of at least 99% of the shortest sequence. These deduplicated protein sequences were then clustered into homologous groups using a three-step iterative process, here again based on different MMseqs2 modules (Steinegger and Soeding 2017). The first clustering step, hereafter called shallow clustering, aimed to gather together very similar proteins. Shallow clusters were built using the *linclust* module with a minimum identity of 90% and a coverage of at least 95% of the shortest sequence. Shallow clusters were subsequently submitted to a second clustering step, called standard clustering. To this end, representative proteins of each shallow cluster were compared and grouped, using the *cluster* module with an e-value of at least 10^-3^, a coverage of 70% of the longest sequence in the alignment, and 30% of identity. Then, standard clusters were grouped into deep clusters using remote homology relationship detection. First, a profile (position-specific scoring matrix) and a consensus sequence were built for each standard cluster. Then, all profiles were compared to all consensus sequences (MMseqs2 with sensitivity -s 4, e-value threshold of 10^-3^, a coverage of at least 60% of the mutual alignment) and the similarities identified were used to group corresponding standard clusters into deep clusters (set-cover clustering). The parameters used in the second and third clustering steps were defined based on results described in the PhaMMseqs pipeline (Gauthier et al. 2022). For more details about the parameters and clustering strategy, see Supplementary Information SI1.

Deep clusters containing at least two deduplicated proteins constitute the final protein groups, hereafter termed enVhogs for *environmental Viral homologous protein groups*. The number of effective sequences (NEFF) was used (Meier and Soeding 2015) to evaluate the sequence diversity inside each enVhog. The higher the NEFF value, the more diverse are proteins in the cluster, and a minimal value of one meaning the proteins of the cluster are identical.

### Estimating the viralness of each enVhog

The occurrence of each enVhog was then quantified in the viral and cellular worlds (Kristensen et al. 2013). To this end, KEGG prokaryotic genomes (version Jan 2018) were first stripped of their prophages, identified using VirSorter (Roux et al. 2015a). Second, the remaining KEGG and RefSeqVirus proteins were separately clustered (MMseqs2, coverage>0.5) and compared to enVhogs (MMseqs2, bit-score>50). All proteins inside a cluster similar to a given enVhog were considered as similar to this enVhog. For each enVhog, the proportion of cells and viruses with at least a protein similar, named Cp and Vp respectively, were computed as follows (see Fig. S1 for detailed examples). First, cellular species are not evenly represented in KEGG, 2068 of the 2612 species having only one genome whereas for example *Chlamydia trachomatis* account for as many as 73 of the 4755 genomes. For each enVhog and each cellular species, the ratio of strains with a match was computed and the Cp was defined as the mean of these ratios. Second, as viruses have multiple independent origins and very variable gene content, two viruses belonging to different viral groups such as classes or order, will only share a limited number of proteins. For example, a protein of a microvirus strain will only be found in other microviruses and the quantitative presence of this protein has to be computed considering only this viral clade. Therefore, RefSeqVirus genomes were first clustered into clades using MCL (inflation of 1.2, weight=percent of shared proteins between virus pairs based on the RefSeqVirus protein clusters), the 18,846 viruses being gathered into 596 clusters and 497 singletons. For each enVhog, only viral clades with at least one protein similar to this enVhog were considered to compute Vp, the other clades being considered as evolutionarily unrelated. Yet, for virus clades only containing one virus (497 singleton viruses), all proteins will automatically be considered as conserved in all the viruses of this clade, and enVhog similar to these proteins will have a Vp of 1. To reduce this bias, a global pseudocount of 24 over all clades was added to the true number of viruses in a clade to avoid giving a too strong weight to these small virus clades. The importance of this pseudocount has been scaled down by the number of clades the enVhog appears in. Formally, the Vp amounts to (see SI 2 for details):

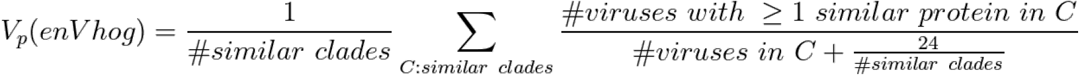

EnVhog viral quotients were computed slightly differently as in (Kristensen et al. 2013):

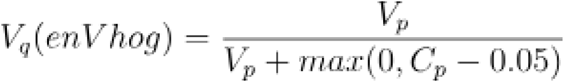

Here, we offset the value of Cp as we realized that some distant (or non-complete) prophages were not detected by VirSorter in KEGG genomes and were still considered part of the cellular world, blurring the signal for proteins scarcely conserved in viral genomes. Manual investigation of specific examples made us tune the Cp offsetting to 0.05.

Considering Vq and Cp, enVhogs were categorized as : (i) ‘Viral’ if conserved in the viral clades they appear in and rarely found in cellular genomes (Vq≥0.8), (ii) ‘Cellular’ if more often conserved in cells than in viruses (Vq<0.5 and Cp>0.05), (iii) ‘Shared’ if conserved by both worlds (0.5<Vq<0.8 and Cp>0.05) and (iv) ‘Unknown’ if not similar to any known viral sequence and not (significantly) conserved in cellular genomes (Vq==0 and Cp<0.05). In addition, viral hallmark annotations were defined by collecting PHROG annotations having the keywords “baseplate”, “head closure”, “head-tail”, “capsid”, “coat”, “head”, “spike”, “tail”, “neck”, “adaptor”, “portal” or “terminase”. After a manual curation step removing more general annotations, 48 annotations were retained (Table S4). All enVhogs matching one of these viral hallmark annotations were added to the ‘Viral’ category.

### Refining enVhog viralness based on the category of contigs on which they appear

The categories describing the viralness of enVhogs are defined above solely on the basis of their similarities to known viral and cellular genomes. However, the nature of the contigs in which they are encoded, currently unknown, could enable their viralness to be determined with greater precision. We thus devised an algorithm that categorizes the nature of contigs from the category of the envhog they encode, and then refine the category of enVhogs using the ones of the contigs they appear in, and so on iteratively (see Fig. S2 for a schematic representation). Briefly, the category (Viral, Shared, Cellular or Unknown) of each enVhog was first transferred to all the proteins constituting them. Then, each contig was tagged as either Viral, Cellular or Unknown if encoding a strict absolute majority of proteins (i.e. if #U > #S+#V+#C in the case of Unknown category, and similarly for other categories, see SI3.a for further details) from the corresponding category, the remaining contigs – which do not support a particular category with strict absolute majority – being tagged as Hybrid. The predicted categories of the contigs are transferred back to the proteins and then used to classify the previously unclassified enVhogs: for each yet unlabeled enVhog (*i.e*. enVhog in category U or S), given its number of occurrences in the different contig categories (V/C/H), a simple Bayesian decision rule (see SI3) decides if there is enough evidence to label the enVhog as Viral or Cellular. When labelled as Cellular, if the Cp (conservation in cellular species, see Materials and Methods) is below 0.05, then the category of the enVhog falls back to Shared class. This strategy (from enVhogs to contigs and back to enVhogs) was run iteratively three times.

### Annotating enVhogs

EnVhogDB clusters were functionally annotated using PHROG (Terzian et al. 2021) and Pfam (Mistry et al. 2021) databases. The moderate size of PHROG allowed us to run HHblits (one iteration, e-values <0.001, and default parameters) in all-against-all comparison between enVhogs and PHROG HMMs. To annotate the enVhogs with Pfam-A profile database (accessed at 22-Oct-2022), we used MMseqs2 *search* with Pfam-A profiles as queries and enVhog representative sequences as target (parameters: sensitivity 5.7, Coverage 0%, E-value 0.001, Coverage mode 0).

### Annotation of proteins from environmental samples using enVhogs

Viral contigs assembled from viromes sampled in four distinct ecosystem types were downloaded, namely sulfuric soils (Bi et al. 2024), grassland soils (Santos-Medellín et al. 2023), surface marine water (Ni et al. 2024) and human gut (Garmaeva et al. 2024). Proteins were predicted using prodigal (parameters -c -m -p meta) and a random set of 1000 proteins was sampled in each of the four sets. These proteins were compared to RefSeqVirus proteins using MMseqs2 (e-value<1e-3). RefSeqVirus proteins were considered as unannotated if they contained the words “hypothetical protein”, “unknown” or whose annotations consisted only of an ORF number. Virome proteins were also compared to PHROG and EnVhogDB HMMs using HHBlits (e-value<1e-3).

To evaluate the results obtained using more sensitive searches against EnVhogDB, the 1,159,978 complete proteins (prodigal; parameters -c -m -p meta) from human gut viral contigs (Zolfo et al. 2024) were grouped into clusters (mmseqs; >70% of bidirectional coverage). For each of the 65,409 clusters, a multiple sequence alignment was computed (MMSeqs2 result2msa) and the corresponding HMM was compared to enVhog HMMs (HHblits; E-value<1e-3). Finally, to test the results obtained using EnVhogDB on a microbiome, the same strategy was used for proteins from a microbiome of IBDMDB (sample PSM6XBTR, arbitrarily chosen; Lloyd-Price et al. 2019), resulting in 41,505 clusters encompassing 75,278 proteins.

## Results

### Leveraging a large number of publicly accessible viral metagenomes

A set of 3,257 publicly available viromes were first identified in 160 different publications. These viromes, sampled from all over the world, were classified as environmental or host-associated and further assigned to 31 environmental subcategories (Fig. 1; Table S1). In the 2,409 host-associated metagenomes, viral communities present in different tissues from a broad range of organisms were sampled, even if datasets from the human gut represent more than half of this category (1,372 viromes). Among the total 12,533,593 contigs (>1.5 Kb), a set of 3,841,422 were detected as viral by VirSorter2, VIBRANT or DeepVirFinder (Fig. S3). As these softwares are based on different approaches (genomic structural features, gene content and/or sequence composition) and different algorithms, a viral origin is more likely to be true if it is predicted by multiple softwares. The number of contigs falsely predicted as viral was minimized by considering as viral only the 1,664,423 contigs predicted by at least two softwares, as in Zayed et al. 2021. Using these contigs, 8,978,898 proteins were predicted.

**Figure 1.**
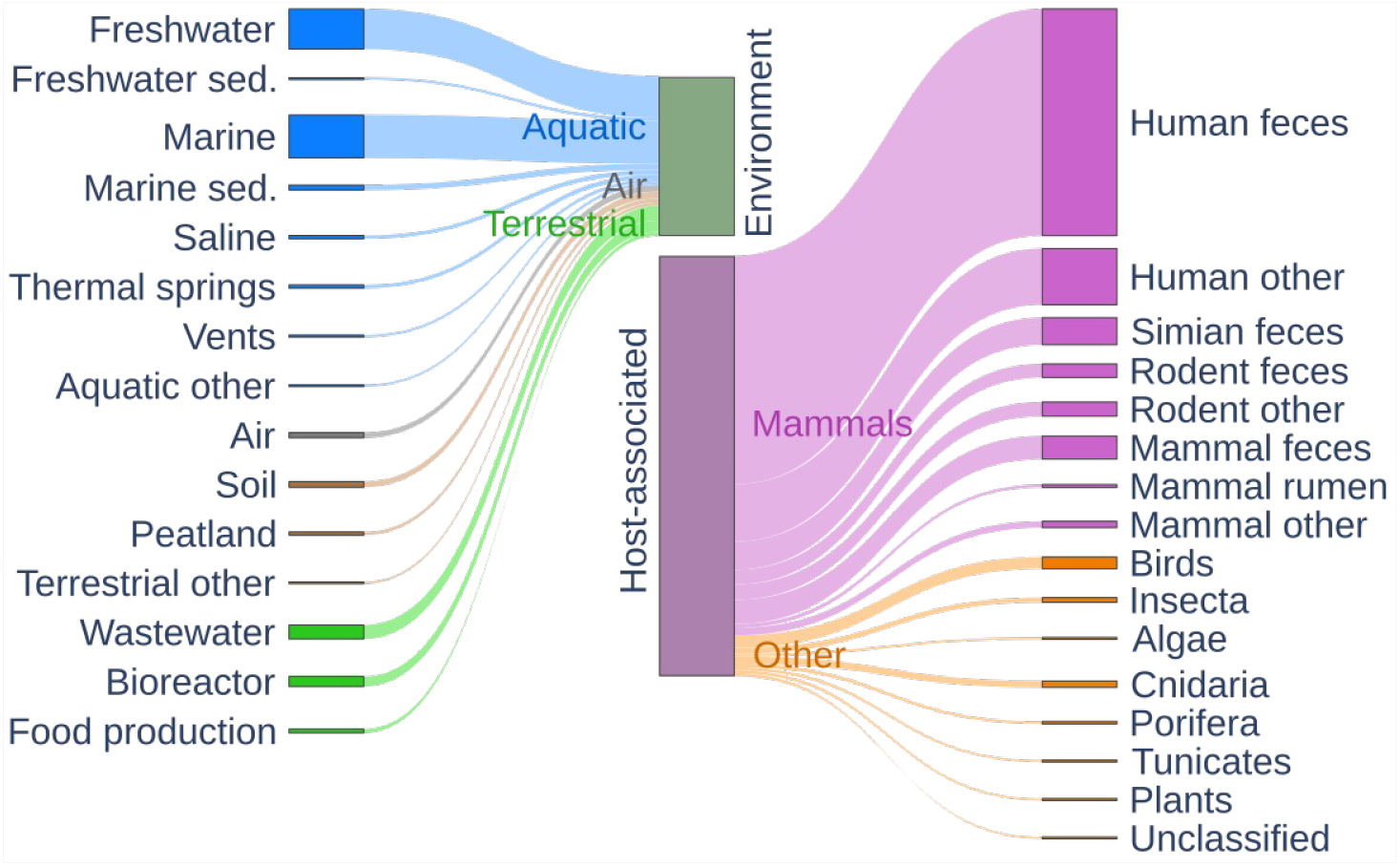
Environmental categories of the viromes assembled for EnVhog. Sankey plot of the ecosystem classification. The 3,257 collected viromes mostly fall into two categories: environmental viromes and host-associated viromes (at the center of the plot). Environmental viromes were further categorized into Aquatic, Air and Terrestrial, whereas host-associated datasets were separated into those associated to Mammals and to Other hosts.

### Creation of the final set of proteins from various sources

In order to create the largest possible set of viral proteins, reference genomes of (pro-)viruses (RefSeqVirus, PHROG) and genomic sequences of uncultured viruses (IMG/VR, GL-UVAB, efam) were added to virome contigs described above. A total of 129.9 M proteins encoded by 7.7 M viral (fragments of) genomes were thus collected (Table S2). Databases compiled in this study are partially overlapping: reference phages are present both in PHROG and RefSeqVirus and identical metagenomes can be found in IMG/VR, GL-UVAB, efam and/or the 3,257 viromes. In addition, viruses with almost identical genomes can be found in RefSeq or in metagenomes from related samples. Protein sequences were thus deduplicated, the resulting 53.6 M distinct sequences being the dataset further described. It has to be noted that the 9 M proteins specifically collected here from public viromes account for 4.4 M deduplicated protein sequences not available in any other viral databases (Fig. 2, left).

**Figure 2.**
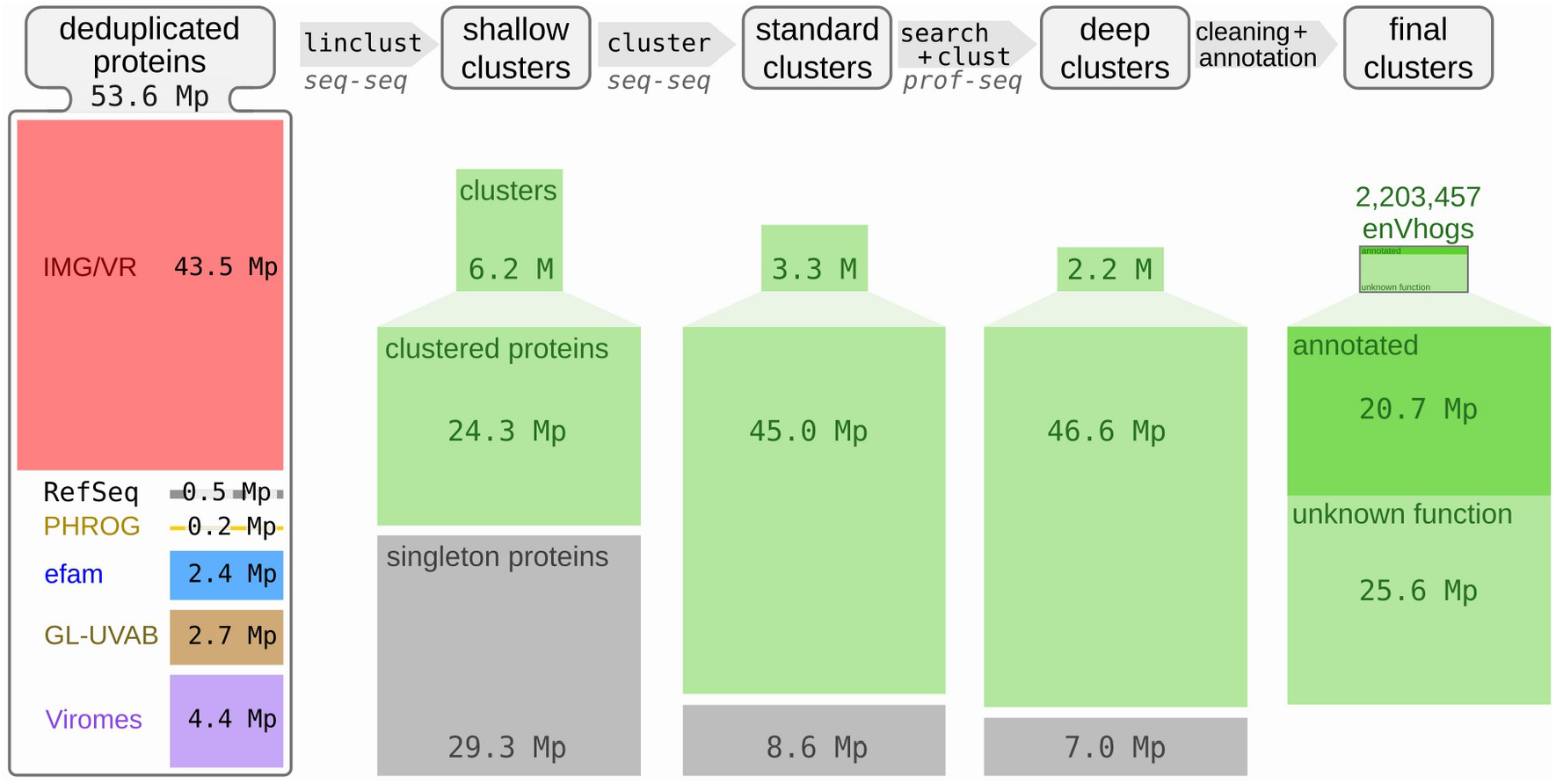
overview of the protein dataset and of the results of the different clustering steps. Left: contribution of each of the data sources in terms of the number of the 53.6 M deduplicated proteins (Mp stands for Millions of proteins). Right : the three clustering steps, realized iteratively (i.e. generated from the results of the previous clustering level) and named shallow, standard and deep clustering and their results in terms of number of clusters and singleton proteins. The number of clustered proteins is also indicated below the number of clusters. At the top, the MMseqs2 modules used, their philosophy and major parameters are indicated (seq-seq and prof-seq stand respectively for sequence-sequence and profile-sequence comparisons).

Considering the taxonomic affiliation, most viral realms are significantly represented, with 0.04 M proteins from *Adnaviria* (archaeal filamentous viruses with A-form dsDNA genomes and alpha-helical MCP), 38.40 M from *Duplodnaviria* (dsDNA viruses with a HK97-fold MCP), 3.43 M from *Varidnaviria*, (dsDNA viruses with a vertical jelly roll MCP), 0.24 M from *Monodnaviria* (ssDNA viruses with a HUH superfamily endonuclease) and 0.48 M Riboviria (RNA viruses with a RNA-dependent RNA polymerase and viruses that encode reverse transcriptase), only *Ribozyviria* (hepatitis delta-like viruses with circular, negative-sense ssRNA genomes) being very rarely found in proteins obtained from metagenomes. The most prevalent viral class is *Caudoviricetes* (38.1 M proteins), the second most abundant class is *Megaviricetes* (2.6 M proteins) (Fig. S4). At the family level, 215 viral families are represented, 138 of which encompass more than a thousand proteins. In addition to these proteins encoded by affiliated contigs, as many as 10.39 M proteins are encoded by contigs that remained unclassified.

### Clustering proteins in three steps that includes profile-sequence comparisons

The 53.6 M proteins were then clustered into homologous groups using a three step procedure. First, 35.5 M shallow clusters were generated, roughly corresponding to groups of proteins encoded by genomes from the same genus, made of 6.2 M clusters of at least two proteins, and 29.3 M singleton proteins (Fig. 2). Secondly, these shallow clusters were grouped into standard clusters using BLASTp-like comparisons detected by MMSeqs2. This methodology is classically used by most databases to generate their protein families, for example in efam (Zayed *et al*. 2021). A total of 11.9 M standard clusters, made of 3.3 M clusters (containing 45 M proteins) and 8.6 M singletons were obtained. Detecting more distant homologous relationships using profile-sequence comparisons, standard clusters were grouped into 9.2 M deep clusters made of 2,210,249 clusters (containing 46.6 M proteins) alongside 7 M singleton proteins. In addition to the total number of clusters decreasing, the average cluster size increases with each clustering step (Fig. S5). Indeed, 50% of the initial 53.6 M deduplicated proteins were grouped in deep clusters containing more than 131 proteins, compared to 42 and 2 for standard and shallow clusters respectively. Deep clusters of at least ten proteins contain more than 75% of the proteins (75.7%), compared to 66.9% for standard clusters. A total of 165 deep clusters larger than 10,000 proteins were generated, and only 91 for standard clusters, two deep clusters exceeding 40,000 proteins (43,904 and 48,424). The 2.2 M deep clusters of more than one protein, representing 87% of the initial set of 53.6 M proteins, will hereafter be termed enVhogs.

To quantify sequence heterogeneity inside enVhogs (*i.e*. deep clusters), NEFF values were computed for each cluster. This sequence diversity measure is the average entropy of all columns constituting a multiple sequence alignment. NEFF values were also computed for PHROGs and Uniclust30 clusters (Mirdita et al. 2016). EnVhogDB clusters were more diverse than UniClust30 homologous groups at every cluster size (Fig. S6) and had a diversity comparable to PHROGs of similar size. For example, for clusters containing from 6 to 20 proteins, the average NEFF values were of 2.5, 2.4 and 2.1 for EnVhogDB, PHROG and Uniclust30 respectively, and of 5, 5.2 and 4 for clusters containing more than 100 proteins. It has to be noted that small clusters (containing between 1 and 5 proteins) were more diverse in EnVhogDB compared to the two other databases (1.44 versus 1.26 and 1.19 for PHROG and Uniclust30 respectively).

### Estimation of the viralness of enVhogs

It has been shown that a fraction of protein families from cultivated viruses are shared by (some) cellular genomes (Kristensen *et al*. 2013). Thus, the viralness of an enVhog, *i.e*. its conservation in the viral world compared to that in the cellular world, is an interesting measure to have for future users. The viralness of each enVhog was estimated through the computation of a viral quotient (Vq) based on its conservation in viral clades it appears in and on its conservation in cellular species. Furthermore, these indices allowed us to categorize each enVhog as being Viral, Shared, Cellular or Unknown. First, 23.2% of enVhogs, containing 56.5% of the proteins, were categorized as Viral thanks to their high Vq (19.7%), their viral hallmark annotation (0.7%) or both (2.8%) (Fig. 3; Table S3; Fig. S7). Only 1.6% of enVhogs (6.2% of proteins) were categorized as Cellular, a quarter of these having no similarity with any known viral sequence (‘No Viral Hit’ column on Fig. 3 and Fig. S7). Similarly, a small proportion of 0.3% of enVhogs (1.5% of proteins) were considered as Shared. Finally, the great majority of enVhogs was classified as Unknown, either because not similar to any sequence (72% of enVhogs, 29.3% of proteins) or not similar to any virus and not significantly conserved in cellular genomes (3.5% of enVhogs, 2.4% of proteins) (Fig. 3).

**Figure 3.**
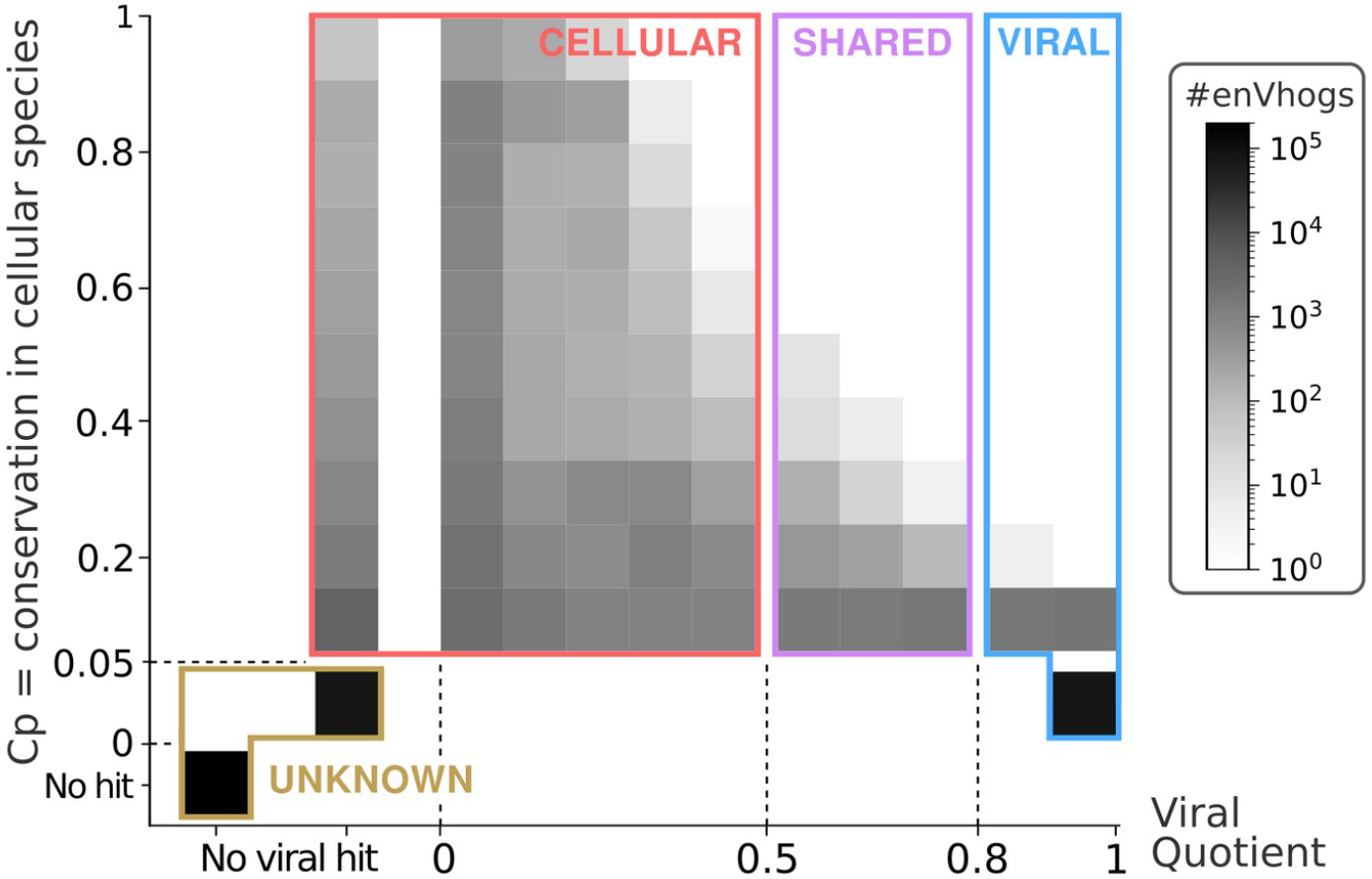
distribution of enVhogs in the four categories (Viral, Shared, Cellular and Unknown). heatmap of the number of enVhogs (colors corresponding to log-scale) according to their conservation in cellular species (Cp on the y-axis) and to their viral quotient (Vq on the x-axis). The categories Viral, Shared, Cellular and Unknown were defined based on thresholds on these Cp and Vq, and are here underlined.

Most enVhogs being categorized as Unknown based on this strategy, we used 3 iterations of our iterative algorithm to transfer back and forth categories from enVhogs to contigs and conversely. As many as 1.5 M enVhogs not similar to a known sequence, viral or cellular, initially categorized as Unknown were re-classified as Viral as they mostly appear on contigs that are finally classified as Viral (Fig. 4; Fig. S8 for details on the first iteration). Viral enVhogs now account for 89% of the whole set (81% of proteins) whereas Unknown enVhogs only account for less than 2% of proteins and enVhogs.

**Figure 4.**
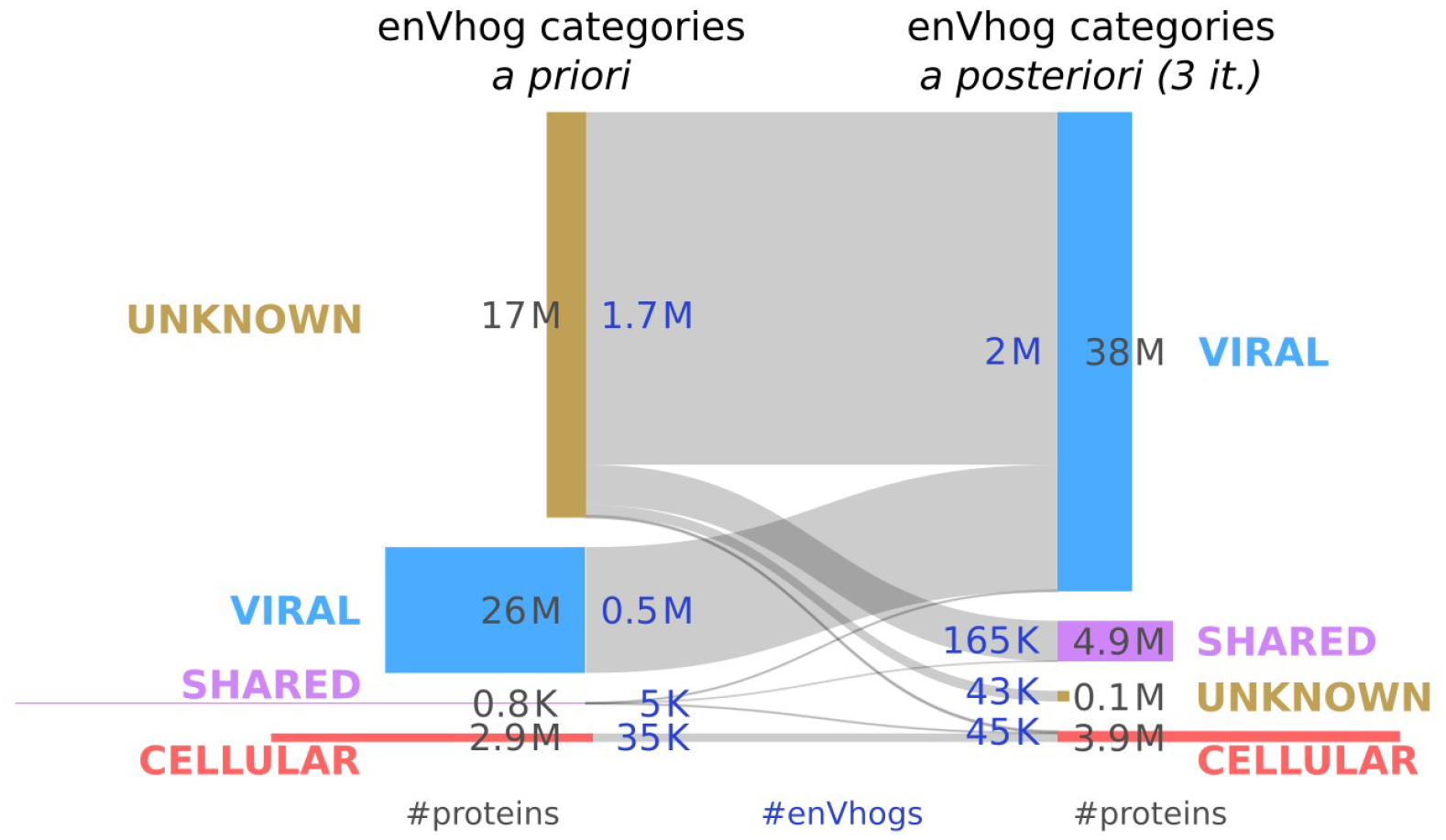
EnVhogDB classes. Sankey plot of the evolution of categories of enVhogs using the iterative procedure. The initial number of enVhogs (in dark blue) and the number of associated deduplicated proteins (in dark grey) in the four categories (Unknown, Viral, Shared and Cellular) are represented on the left. The same numbers are displayed after three iterations of the procedure (on the right). Height of boxes is proportional to the corresponding number of enVhogs and their surface to the number of proteins.

Although its aim was to provide users with a measure of the viralness of each enVhog, the procedure described above also predicted the origin of contigs. In addition to ascertaining the viral origin of most contigs, a limited set of 109,959 contigs were predicted to be of cellular origin. Despite the strict criteria used to identify viral contigs and our use of well-established data sources, it is not surprising to find a small percentage (1.43%) of contigs predicted to be of cellular origin by a complementary approach such as the procedure developed here. Some contigs from all databases were predicted as Cellular (66,541, 36,359, 5,434, 1,516 and 109 in viromes, IMG/VR, eFAM, GLUVAB and PHROG respectively). To improve the quality of our database, we decided to remove (i) these contigs, (ii) the proteins they encode and (iii) the 6,792 enVhogs having lost more than half of their deduplicated proteins and those which became too small (containing less than two proteins). Unsurprisingly, the filtered out enVhogs are enriched with hits against PFAM (35.5%) and cellular proteins (64%) compared to the rest of enVhogs (7.9% and 14.7% respectively). The remaining 2,203,457 enVhogs constitute the final set of EnVhogDB and are simply referred to as enVhogs hereafter.

### Functional annotation of enVhogs

To provide a valuable viral protein resource, enVhogs were annotated using multiple databases and remote homology detection tools. A total of 292,089 enVhogs were similar (HHblits 1 iteration, eval<0.001) to at least one annotated PHROG (Terzian *et al*. 2021), leading to the annotation of 13.3% of enVhogs encompassing 39.9% of clustered proteins (18.5M). These results are very much alike those obtained for PHROGs, for which only 13.1% protein clusters are annotated yet representing a large proportion of proteins (50.6%). An additional 245,777 enVhogs were similar to unannotated PHROGs (23.3% of proteins). Among the ten large functional categories of PHROG, the most abundant functions were from the categories “Tail”, “DNA, RNA and nucleotide metabolism” and “Head and packaging” (see Table 2 for details). A total of 175,246 enVhogs were similar to Pfam-A profiles (13.7 M proteins). Out of these, 32,827 concerned enVhogs that had no hit in PHROGs. Leaving aside similarities to PHROGs or PFAMs of unknown function, 349,938 enVhogs were annotated (15.9% of enVhogs containing 44.8 % of proteins).

**Table 2.** EnVhog annotation according to similarities to PHROG. Total number of annotated enVhogs and proteins for the ten PhROG functional categories.

| Annotation classification | #enVhogs | #proteins |
| --- | --- | --- |
| unknown function | 245,777 | 10,787,649 |
| tail | 78,210 | 3,284,209 |
| DNA, RNA and nucleotide metabolism | 65,269 | 4,621,857 |
| head and packaging | 51,848 | 4,563,053 |
| other | 26,931 | 2,005,222 |
| transcription regulation | 22,091 | 758,551 |
| moron, auxiliary metabolic gene and host takeover | 14,949 | 949,671 |
| connector | 12,148 | 1,120,256 |
| lysis | 12,128 | 785,524 |
| integration and excision | 8,514 | 376,582 |

### EnVhogDB improves virome annotation

To quantify EnvhogDB’s contribution to annotating new viral proteins, proteins from recent viromes from four different ecosystems were annotated using three databases: RefSeqVirus (sequence-sequence comparisons), PHROG and EnVhogDB (sequence-HMM comparisons). The use of EnVhogDB doubled both the proportion of sequences with identified homologs and the proportion of functionally annotated proteins, compared with the reference databases RefSeqVirus and PHROG. EnVhogDB indeed provided functional annotation for 40% to 56% of the proteins in the four new viromes, while RefSeqVirus and PHROG annotated only 19% to 36% and 23% to 35% of the same proteins respectively (Fig. 5A). While EnVhogDB’s improvement over other DBs is the same in every ecosystem, viruses from some ecosystems are clearly better characterized than others. Unsurprisingly, more than 80% of proteins of viral communities from the human gut and from marine surface waters have homologs in EnVhogDB, half of the whole protein datasets being annotated (respectively 56 and 51%). In comparison, a particular, little-studied extreme ecosystem like sulfuric soils with a pH lower than three, has only 69% of its proteins characterized and 40% annotated.

**Figure 5.**
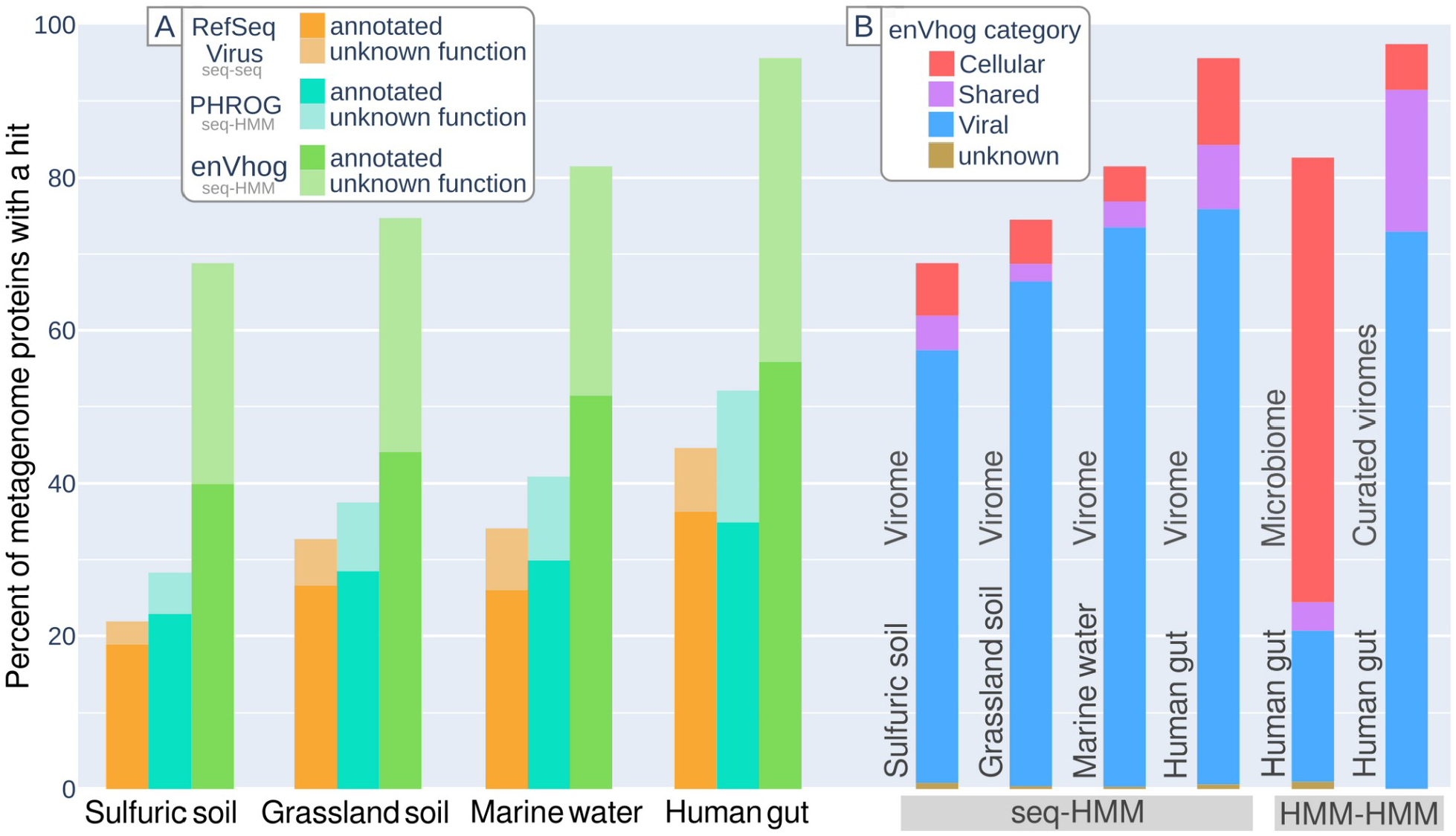
A: Functional annotation of virome. Percent of characterized and annotated proteins from four sets of contigs from different ecosystems using different databases (RefSeq Virus, PHROG, EnVhogDB). B: Categorical annotation of metagenomic proteins (V/SC/U). Protein category (Viral/Shared/Cellular/Unknown) breakdown as identified by hits against enVhog on the four selected viromes, a microbiome and a curated set of human gut viruses extracted from metagenomes.

Based on the same comparison results, the proportion of proteins similar to enVhogs of the four defined categories (Viral, Shared, Cellular, Unknown) was calculated for each of the four viromes (Fig. 5B). Results are comparable for the four viromes : a negligible proportion of Unknown enVhogs, a dominance of Viral enVhogs (>78%), the remainder being made up of Cellular and Shared enVhogs. While this pattern is expected for viromes, results are very different when applying the same analysis to a human gut microbiome (Lloyd-Price et al. 2019). First, most proteins from the human gut microbiome are similar to an enVhog (82%) and in more than 70% of the cases, it concerned enVhogs of the Cellular category. Given that EnVhogDB have been cleaned from enVhogs only supported by Cellular contigs, this result suggests that most of the proteins encoded by microbes in the human digestive tract have an homologous counterpart in the viral world. Second, the proportion of proteins detected as Viral amounts to 20% of the microbiome. As seen previously, almost all human gut virome proteins are characterized and thus the uncharacterized proteins (18%) of the microbiome are probably mostly of cellular-only origin.

It should be noted that for the annotation of the latter dataset, its proteins were first clustered (75,278 proteins have been clustered into 10,391 clusters and 31,114 singletons), with the resulting cluster HMMs being compared to the EnVhogDB HMMs. The same methodology was then applied to a set of human intestinal viral genomes (Zolfo et al. 2024). Thanks to this more sensitive strategy, a homolog in EnVhogDB could be identified for over 97% of the proteins in these genomes.

## Discussion

### Moving closer to a comprehensive view of viral genomic sequence space

First, we collected as many viral protein sequences as possible. In addition to existing viral databases mostly built from sequences extracted from microbial genomes and microbiomes, an effort was made here to identify, collect and assemble thousands of published viromes whose contigs are absent from viral databases. As many as 4.4 million “new” proteins were added by these datasets, sampled from a wide range of ecosystems and coming from all over the world. This contribution of more than seven times the number of proteins of reference viruses present in RefSeqVirus is therefore important for continuing to move closer to a comprehensive view of viral protein sequence space. Yet, the EnVhog database may still suffer from different biases. First, the collected samples, both for viromes or for microbiomes, are usually biased toward anthropocentric environments. Indeed, a larger proportion of protein were characterized in human gut or marine systems compared to soils or extreme environments. This results in a biased representation of the viral diversity on Earth. Second, the biochemistry processes are usually focused on DNA molecules (especially double stranded), RNA viruses being poorly represented in the available raw data. Third, to minimize the presence of contigs of cellular origin, only contigs predicted as viral by two detection softwares were considered. However, many (fragments of) viral genomes may have been omitted by these tools, especially viruses that are only distantly related to reference viruses. The enVhogs generated here, more representative of the overall viral diversity, could prove useful for revisiting these datasets to identify more *bona fide* viral contigs.

### EnVhogDB : a large set of diverse families

To use these millions of proteins efficiently, a key step is to organize them into families of homologous proteins based on sensitive similarity search tools. Existing databases that include metagenomic data provide either individual proteins (IMG/VR) or protein cluster HMMs (eFAM), the latter being generated by standard BLASTp-like comparisons. Our three-step clustering procedure, the last step of which uses profile-sequence comparisons, provides better clustering than standard methods (Gauthier et al. 2022). This result illustrates once again that the diversity of viral families requires sensitive similarity search methods. The HMM-HMM comparisons used to cluster reference proteins in PHROG are even more sensitive (Terzian et al. 2021), and the detection of even more distantly related homology would make it possible to merge certain enVhogs. Yet, the computational greediness of such comparisons ruled us out of their application on this protein set and the number of viral protein families contained in this dataset is thus overestimated. Yet, counter-intuitively, the internal diversity of enVhogs is comparable to that of PHROGs. This is probably due to the nature of the genome sampling making up these two DBs:

(i) PHROG reference viruses, isolated from a small number of hosts and ecosystems, represent a highly fragmented view of the viral sequence space, and despite the sensitivity of HMMs, linking parts of this space is not possible in the absence of stepping stones, (ii) EnVhogDB metagenomic contigs are recovered at random from very diverse ecosystems and therefore better represent the viral genomic space, making it easier to group the encoded proteins. As a result, enVhogs are both as internally diverse and much more numerous (2M instead of 40K) than existing viral homologous groups.

### A majority of new viral proteins can be characterized by EnVhogDB

Their large number and diversity make the detection of similarities between enVhogs and new viral proteins more efficient than using databases built solely from reference viruses. Indeed, the use of EnVhogDB significantly increased the proportion of proteins with a homolog in new genomes (now more than two thirds). Even if a non-negligible but difficult-to-estimate proportion of viral diversity remains to be discovered, EnVhogDB contains a large proportion of the protein families encoded by viruses that are currently recognizable. In addition, its use improved by 54%-74% the proportion of annotated proteins in new genomes, which can be explained in two complementary ways. Firstly, the enVhogs themselves were annotated in depth using sensitive homology detection techniques (i.e. HMM-HMM comparison with PHROGs and PFAMs). Secondly, the 350K annotated enVhogs, made of 20.7M proteins, are internally diverse and well adapted for detecting distant homologies between new proteins and enVhogs (sequence-HMM). An even greater number of new proteins could be annotated when these proteins were themselves organized into homologous groups and compared with enVhogs using HMM-HMM tools.

Functional annotation is crucial to the description and understanding of viruses through their genomes, and even if EnVhogDB represents a major step forward in identifying function encoded in viral proteins, a large fraction of enVhogs remains of unknown function. The functions of these enVhogs will have to be elucidated experimentally or using three-dimensional structure predictions and comparisons, a strategy complementary to our sequence-based approach.

### Quantifying the viralness of these clusters, even of previously unseen ones

For each protein family, we quantified its conservation in the viral world compared to the cellular world, allowing us to assess the viral origin of about a quarter of the protein clusters. Yet, a majority of enVhogs had no similarity to known viruses, illustrating that reference viruses are far from representing an exhaustive view of the viral world. Then, these categories were refined by an iterative cluster-to-contig-to-cluster strategy. First, this strategy allowed us to identify and remove the small proportion of contigs and clusters whose viral origin was unclear and thus provide a database as clean as possible. Second, this strategy reduced the number of clusters initially classified as Unknown from 75% to 2%, confirming our view that reference viral genomes are poorly representative of real diversity in the environment. In addition, 89% of the EnVhogDB is constituted by proteins for which we are confident about their viral origin. EnVhogDB clusters classified as Cellular or Shared should not be removed as they represent families more or equally conserved in the cellular than in the viral world (Kristensen et al. 2013). These contain, among other things, auxiliary metabolic genes, and it would be interesting to analyze the extent of the functional diversity occasionally present in viruses on the basis of this dataset. Indeed, most microbiome proteins were similar to Cellular enVhogs, suggesting that most of the proteins encoded by microbes in the human digestive tract have at some point been carried by viral genomes. These categories also highlighted that 20% of the human gut microbiome proteins were of viral origin, a proportion likely to be realistic thanks to EnVhogDB completeness. This proportion, markedly high, could be explained by (i) the significant proportion of prophages in microbial genomes (Roux et al. 2015a), a proportion still probably underestimated, (ii) the presence of free viral particles, both phages and eukaryotic viruses, captured and sequenced in microbiomes (Schulz et al. 2020), (iii) the presence of cells infected by viruses in their lytic cycle whose genome number of copies ease them to pass the detection threshold (López-García et al. 2023). The categories defined here to describe the viralness of clusters are thus useful for discriminating typical cellular and viral proteins, and will help quantify the presence of cellular DNA sequences in metagenomes and or identify of (pro)viruses in cellular (meta)genomes.

## Conclusion

The EnVhog database is currently the most up-to-date ressource for viral protein families, covering a very broad spectrum of environments and geographic locations on Earth. EnvhogDB significantly extends previous viral protein databases in terms of diversity and completeness, and provides highly sensitive HMMs for protein annotation. Through this work, we hope to provide a dataset that will help the microbiology and ecology community to identify new viruses and better understand the functioning of newly detected viral genomes.

## Data availability

EnVhogDB clusters as MSA and HMMs, as well as profile database (MMseqs2 format), are available online for user’s download at http://envhog.u-ga.fr/envhog/

In addition, information on contigs, proteins, intermediate clusters and enVhogs is available on the same website via various tables in tsv format (details on file format and headers are given at http://envhog.u-ga.fr/envhog/) :

> *envhog_infos.tsv*: for each EnVhogDB cluster, a description of size, maximum protein length, diversity, functional annotations, viralness, hostess, viral quotient and category V/S/C/U as identified by our iterative algorithm. *prot_dedup_shallow_standard_cluster_envhog_contig.tsv*: for each original protein, a list of its : (i) the original protein, (ii) the deduplicated protein group, shallow, standard and deep clusters and enVhog to which it belongs and (iv) its original contig.
>
> *enVhog_classes.tsv.gz*: for each enVhog, its categories V/S/C/U at each iteration of our iterative algorithm
>
> *contigs_infos_class.tsv*: for each contig, the contig identifier, its accession number in the original dataset, figures on the contig (length, number of proteins) and information on its taxonomy and number of enVhogs breakdown by categories V/S/C/U, for iterations 0 and 3 of our iterative algorithm
>
> *enVhog_contigs_classes.tsv.gz*: Category V/C/H/U for each contig at each iteration

The scripts used to create EnVhogDB are available on the git repository https://gricad-gitlab.univ-grenoble-alpes.fr/galiezc/envhog

## Acknowledgements

The computations presented in this paper were performed using the GRICAD infrastructure (https://gricad.univ-grenoble-alpes.fr), which is supported by Grenoble research communities, and using the Froggy platform of the GRICAD infrastructure (https://gricad.univ-grenoble-alpes.fr), supported by the Rhône-Alpes region (GRANT CPER07_13 CIRA) and the Equip@Meso project (reference ANR-10-EQPX-29-01) of the programme Investissements d’Avenir supervised by the Agence Nationale pour la Recherche.

Preprint version 3 of this article has been peer-reviewed and recommended by Peer Community In Microbiology (https://doi.org/10.24072/pci.microbiol.100152). We would like to sincerely thank the reviewers and our recommander Craig Herbold for helping us improve our manuscript and supplementray data.

## Funding

This work was developed in the framework of Grenoble Alpes Data Institute, supported by the French National Research Agency under the “Investissements d’avenir” program (ANR-15-IDEX-02). This work was supported by the European Research Council’s Horizon 2020 Framework Programme for Research and Innovation (“Virus-X”, project no. 685778).

## Conflict of interest discolsure

The authors declare that they comply with the PCI rule of having no financial conflicts of interest in relation to the content of the article.

## Supplementary materials

## Supplementary tables

Table S1. Description of the 3,257 collected viromes.

**Table S2. Dataset summary:** Almost 130 million proteins were selected from multiple resources. All proteins were cleaned to remove sequences with intermediate stop codons as well as sequences with 5 or more unknown amino acids.

**Table S3**. Numbers and proportions of enVhogs and proteins with different thresholds of Vq index.

**Table S4**. List of the 48 hallmark annotation terms defined.

Supplementary figures and supplementary information can be found after the references

## Supplementary materials

## Supplementary figures

**Figure S1.**
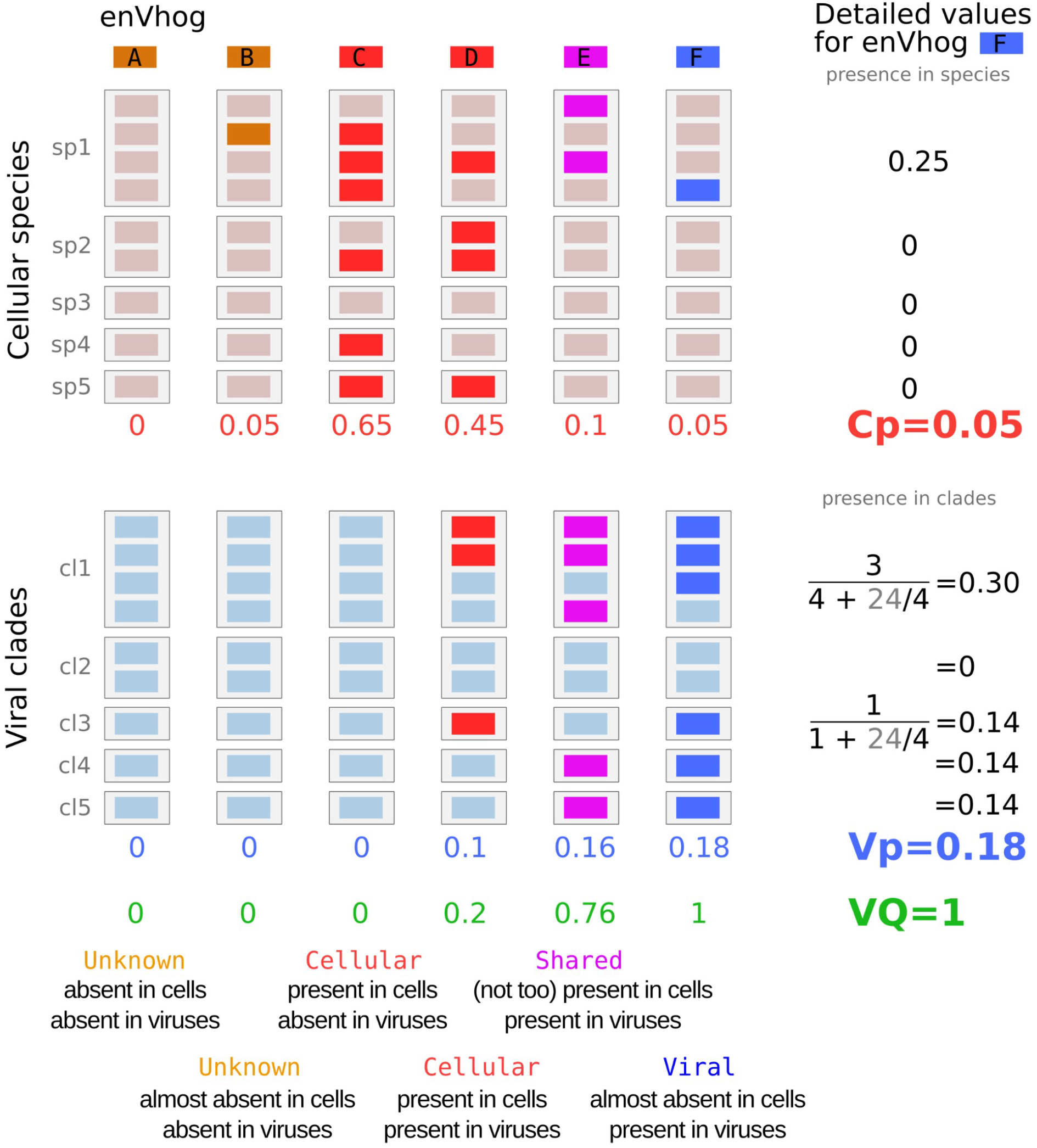
Example of the categorization as Viral, Shared, Cellular or Unknown for six enVhogs (named A to F). For each enVhog, displayed as columns, its presence in cellular species (sp1 to sp5) are first determined using similarity searches and are further used to define a measure of Cellular presence, or Cp. Numbers and formula are detailed for the enVhog on the right (enVhog F) whose Cp is 0.05. Then, the presence of each enVhog in viral clades (cl1 to cl5) are determined (MMseqs2) and are used to compute a measure of Viral presence, Vp. The Vp of enVhog F is 0.18. Cp and Vp are then used to compute the Viral Quotient (VQ).

**Figure S2.**
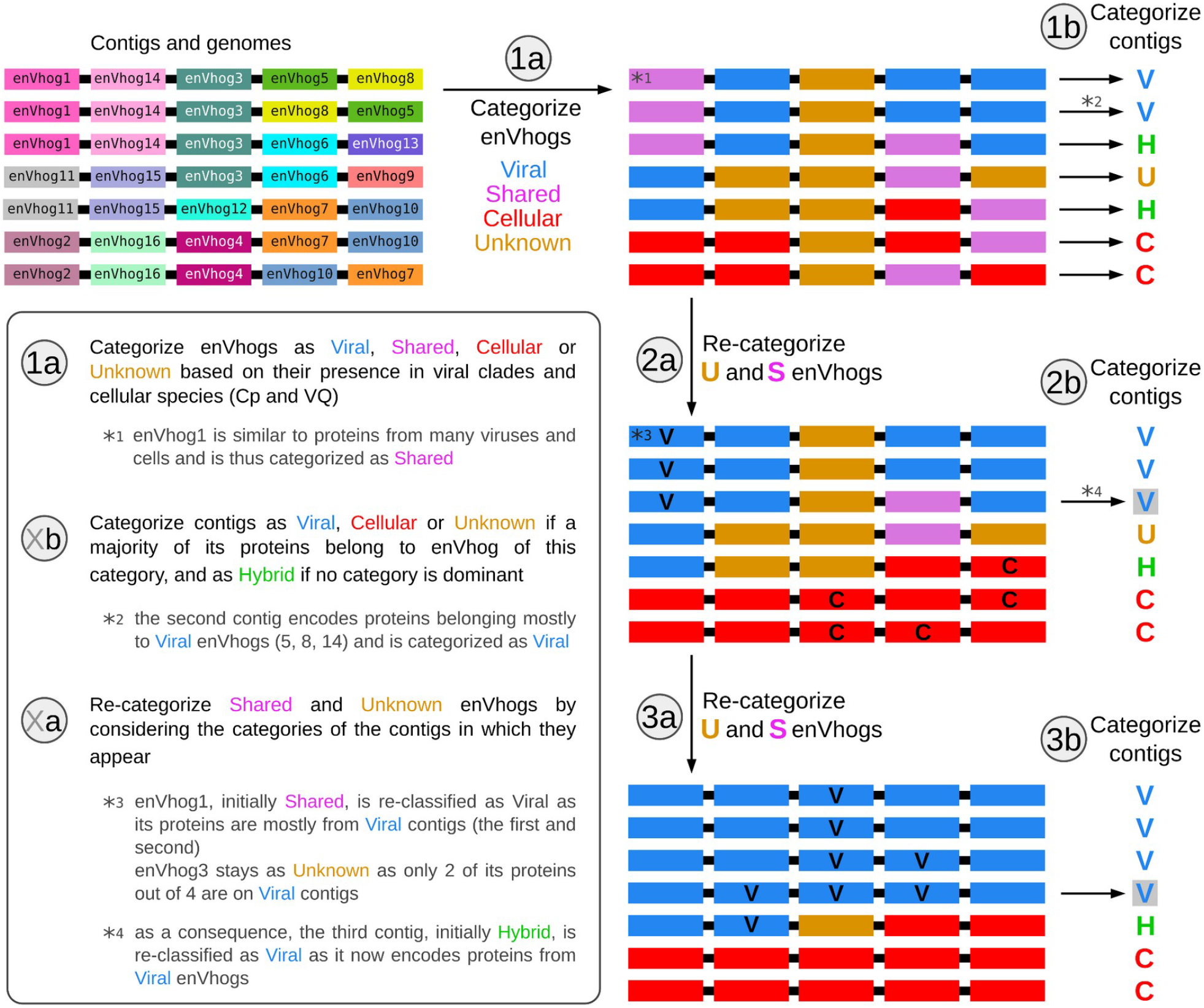
schematic explanation of the iterative procedure used to define the category of each enVhog based on the category of the contigs in which they are encoded. Seven contigs are represented, each encoding four proteins that were clustered into 13 enVhogs displayed in colors at the top left. Each enVhog is categorized independently using Cp and Vq values (step 1a) and then, contigs are classified based on the most abundant category of the enVhog they encode (step 1b). The category of enVhogs was then refined using the category of the contigs on which they appear, and so on (steps Xa and Xb).

**Figure S3.**
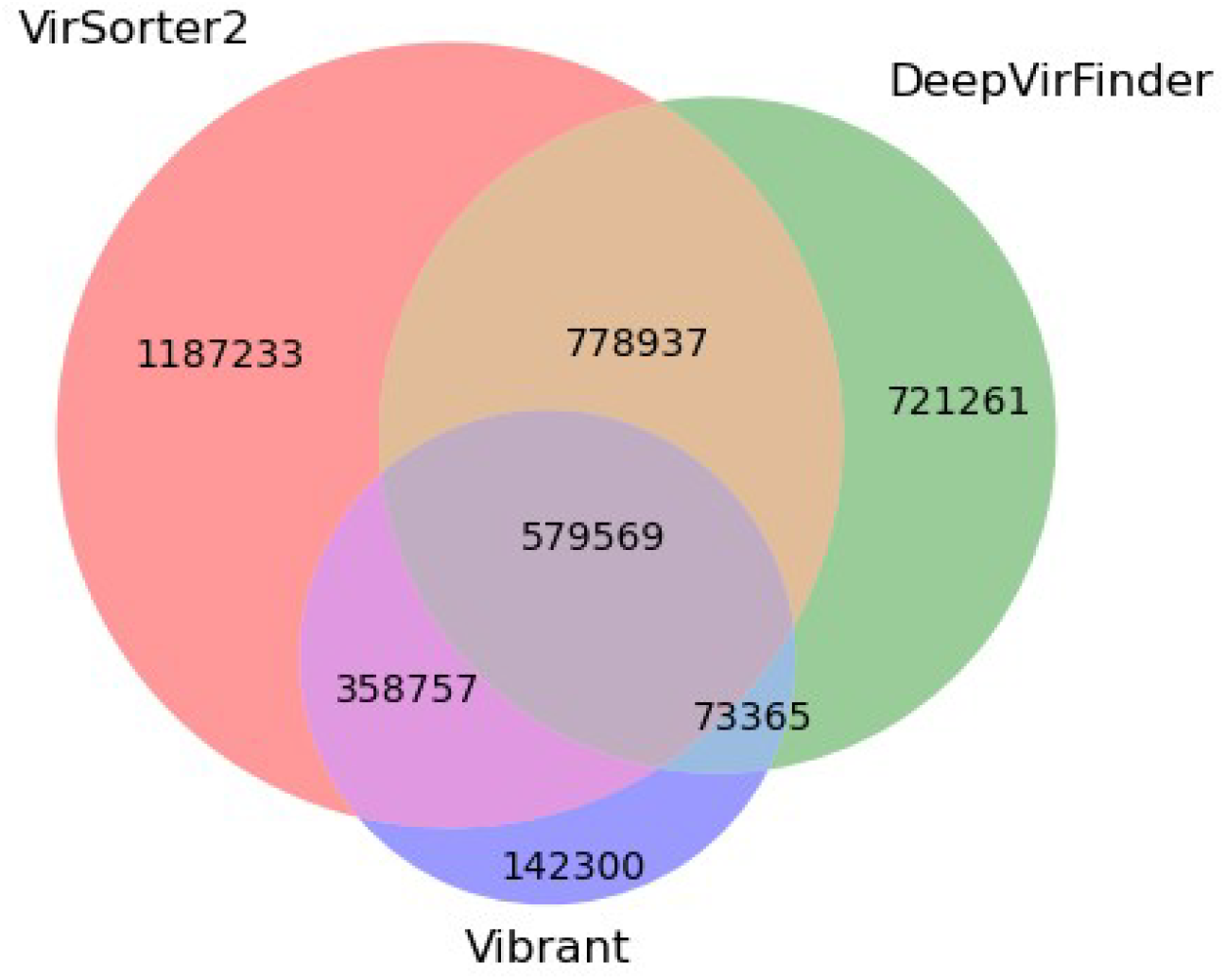
Venn diagram of the viral contig prediction of our manually assembled contigs by VirSorter2, Vibrant and DeepVirFinder. Contigs kept in EnVhogDB lie at the intersection of at least 2 out 3.

**Figure S4.**
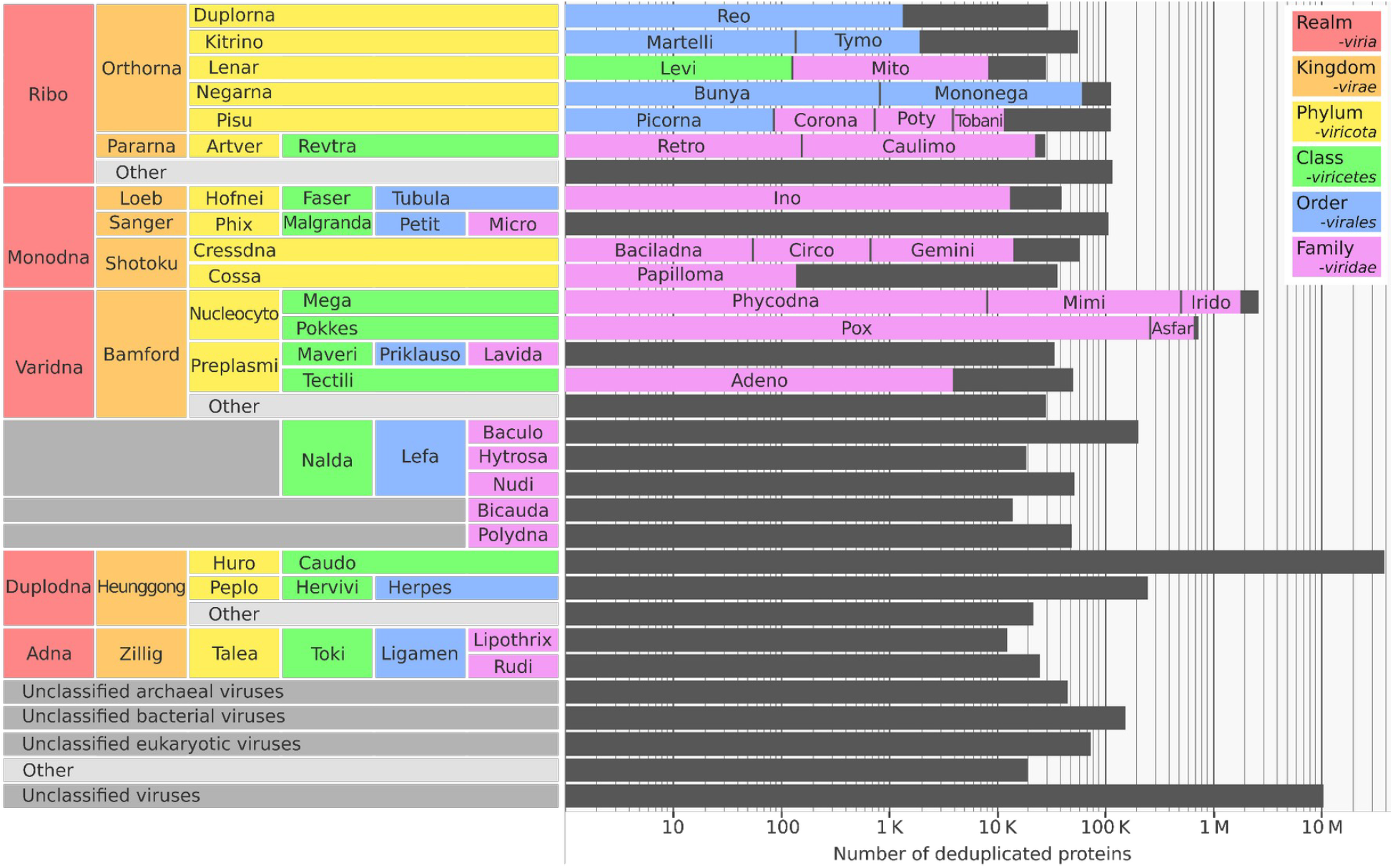
Taxonomic affiliation of the 53 million deduplicated proteins.

**Figure S5.**
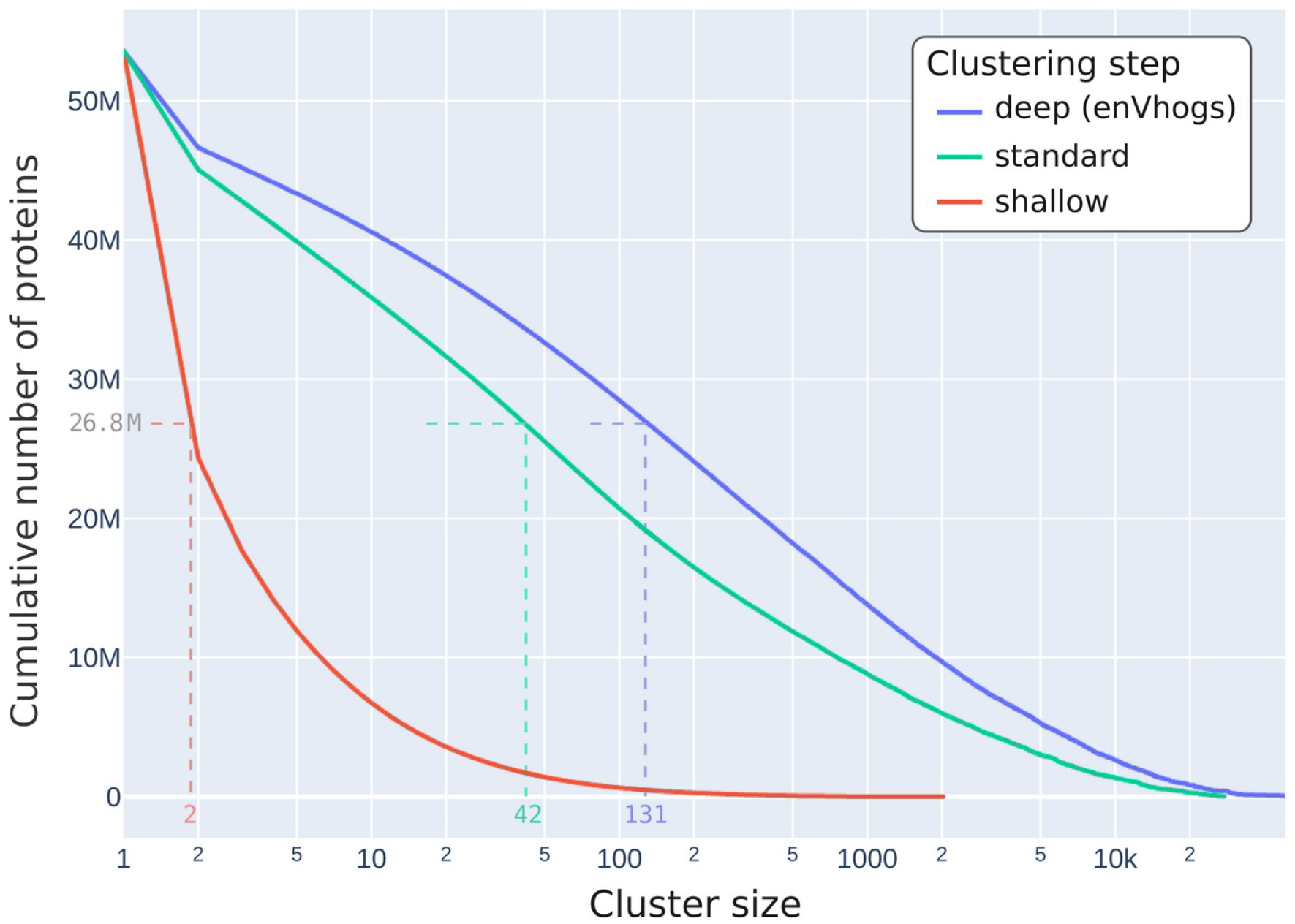
Clustering structure. The 53.6 million protein sequences were grouped into enVhogs using a three step clustering procedure. The cumulative number of proteins per cluster size are displayed for clusters at each step, named shallow, standard and deep clusters, these last clusters corresponding to enVhogs and singletons. Half of the proteins are grouped respectively in clusters containing more than 2, 42 and 131 sequences.

**Figure S6.**
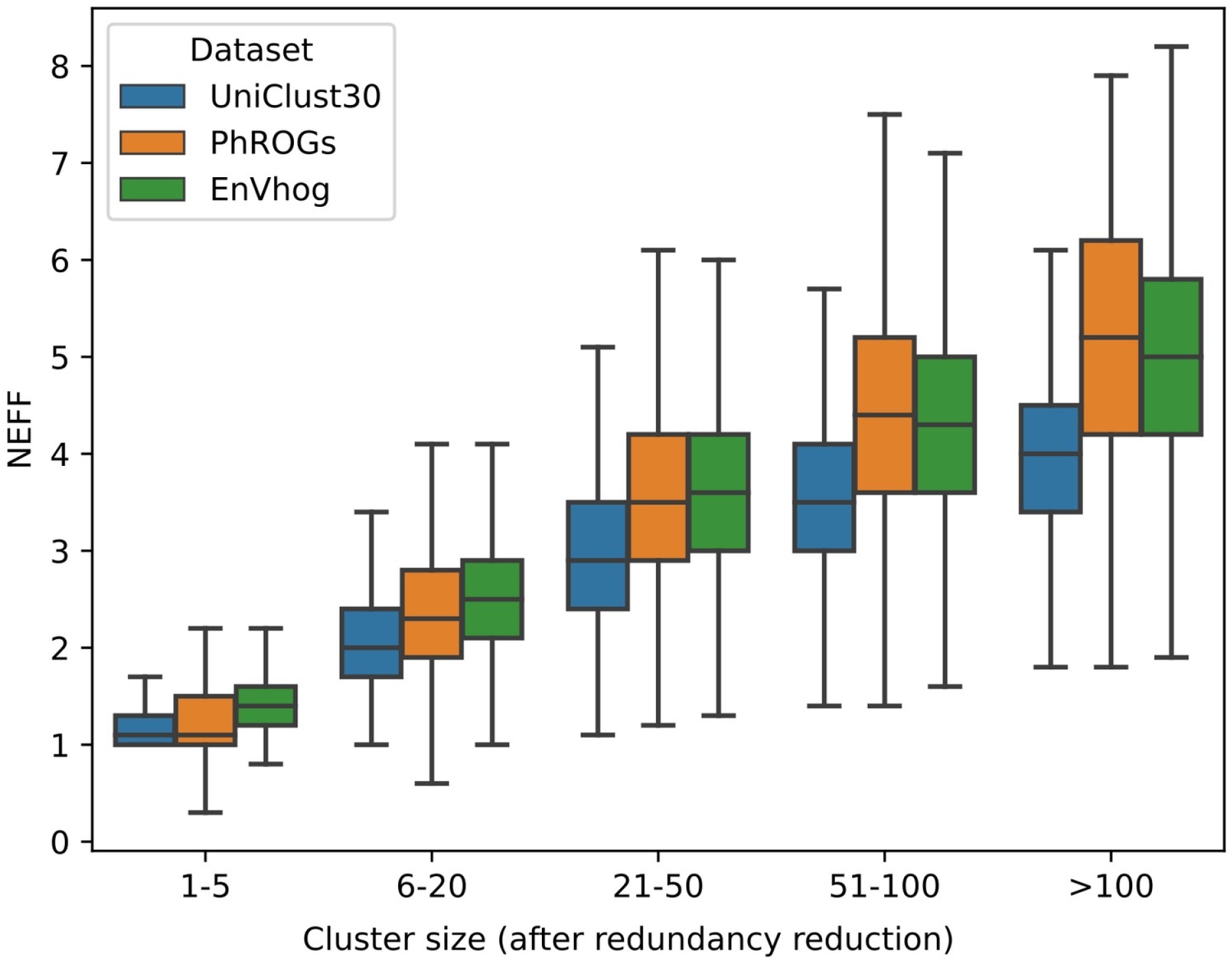
Sequence diversity in EnVhogDB clusters: NEFF values were calculated for all enVhogs and compared to the one of protein families from the reference databases PHROG and UniClust30. On the x-axis, the categories correspond to the number of sequences in the cluster (as computed by HHblits after redundancy reduction).

**Figure S7.**
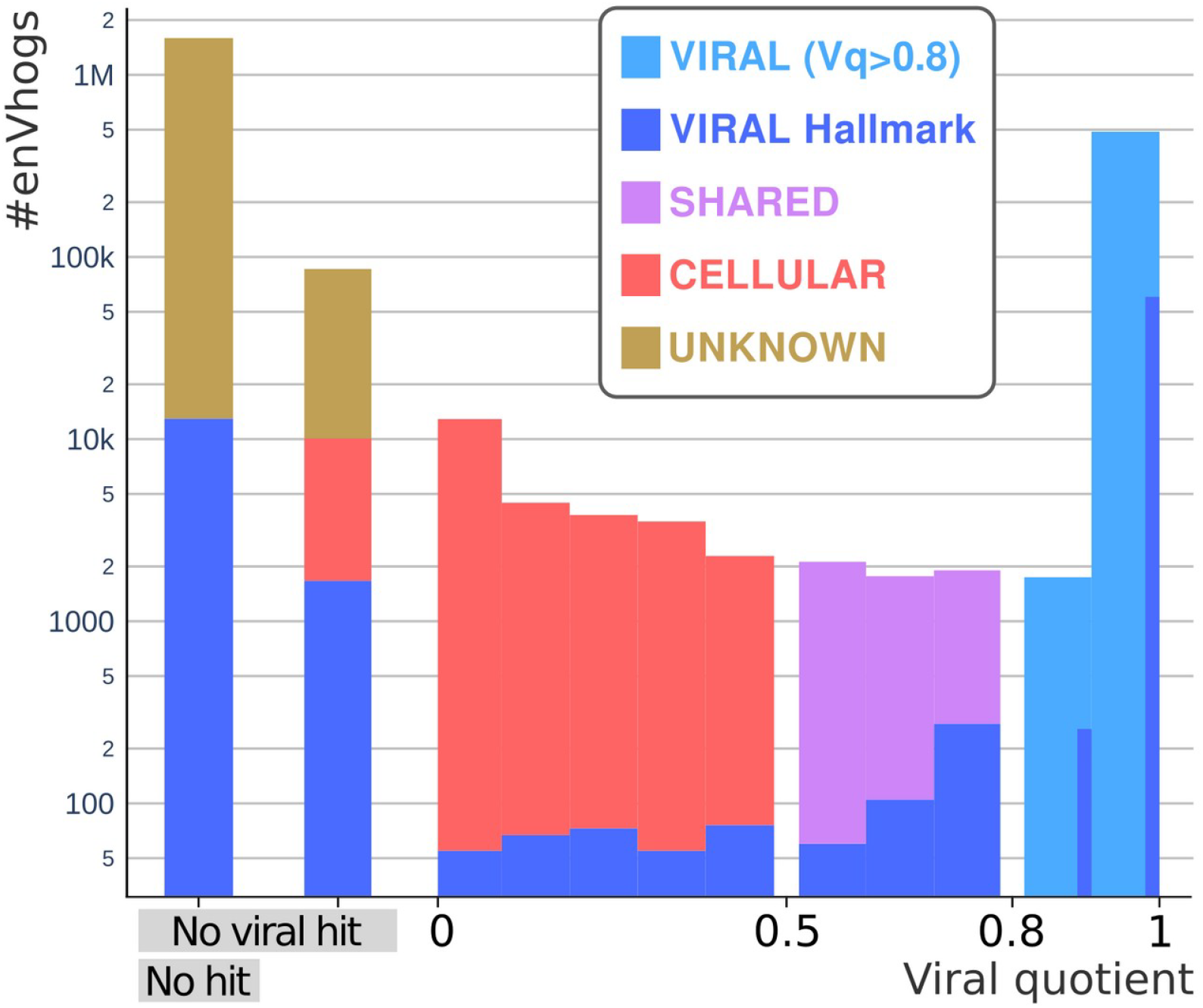
distribution of enVhogs in the four categories (Viral, Shared, Cellular and Unknown). Number of enVhogs in the four categories (Y-axis, in log-scale) according to their Viral quotient (Vq) in the x-axis. Bars on the left correspond to enVhogs with no hit in viral and cellular databases and no hit against the viral protein database. EnVhogs are classified as Viral if their Vq is greater than 0.8 or if they were identified as viral hallmarks.

**Figure S8.**
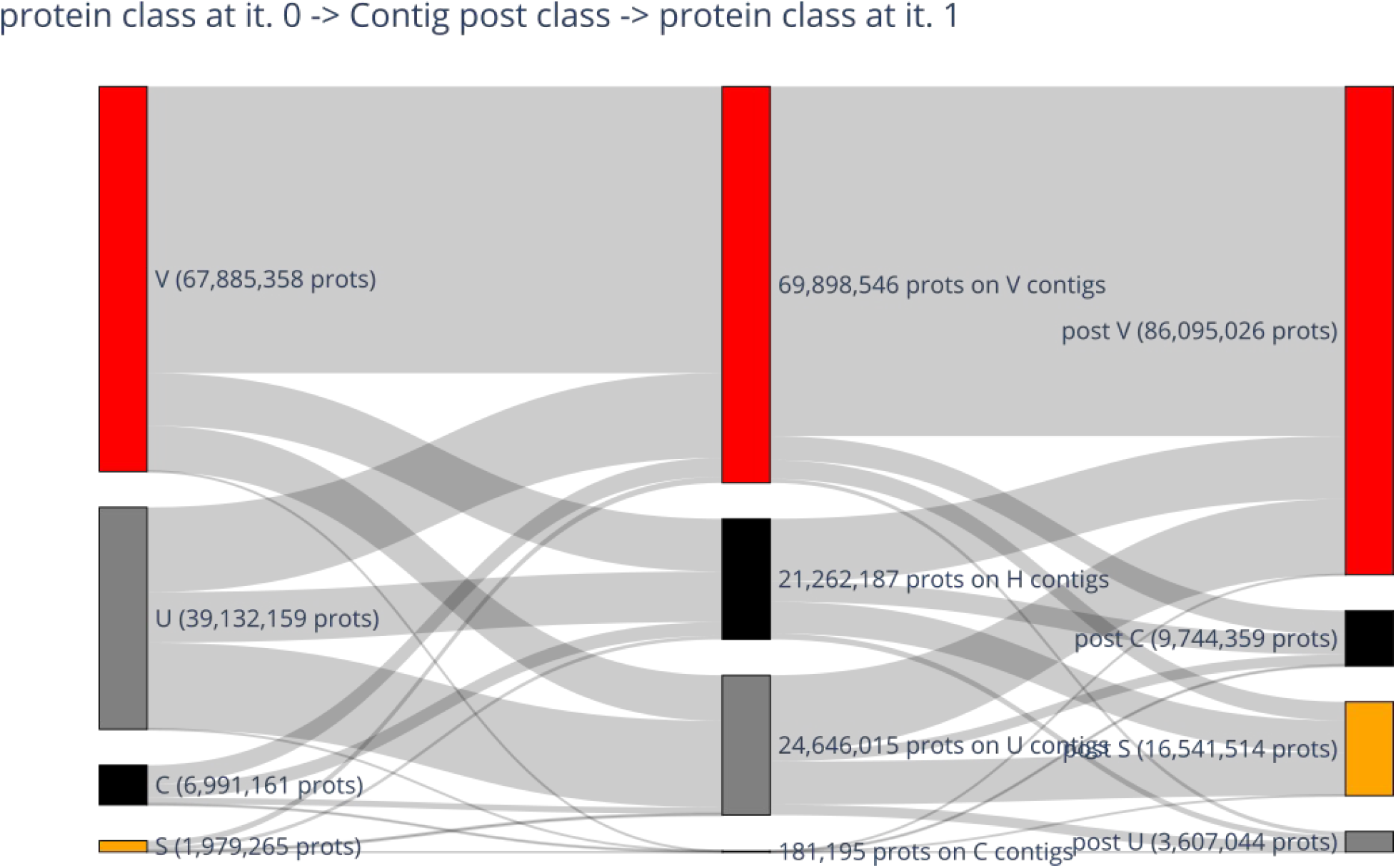
Evolution of cluster classes in our iterative algorithm taking advantage of genomic context. Left column: Original cluster class, center column: Posterior contig class (V/B/U), rightmost column: Posterior cluster class after 1 iteration. With a single iteration, most of the proteins belonging to enVhogs of category U got an assigned category in V/C/S. To generate the final classification 3 iterations of this algorithm have been run.

## Supplementary Table

**Table S4.**
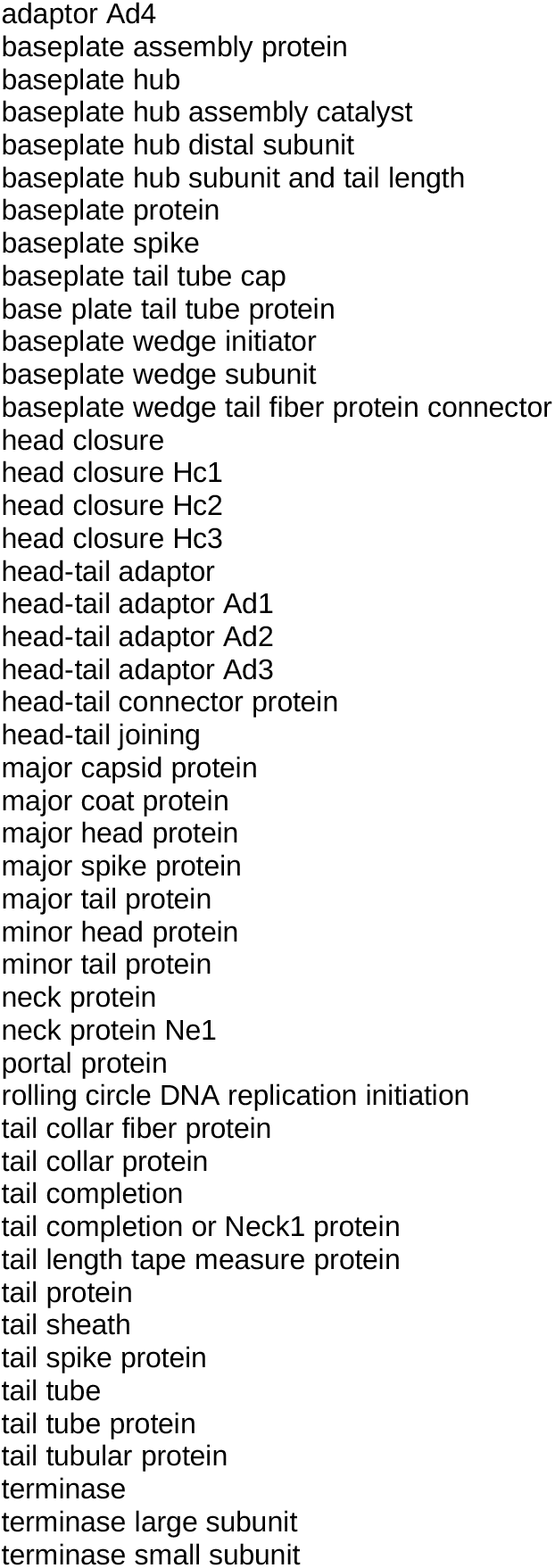
List of the 48 hallmark annotation terms defined.

## Supplementary information

### SI1: Similarity search, clustering and HMM generation parameters

Redundancy reduction step (i.e. deduplication):

~~~
Linclust parameters: -c 0.99 --cov-mode 1 --seq-id-mode 1 --cluster-mode 2 --min-
seq-id 0.99 -e 0.001 --kmer-per-seq 100
~~~

Shallow clustering:

~~~
Linclust parameters: -c 0.95 --cov-mode 1 --seq-id-mode 1 --cluster-mode 2 --min-
seq-id 0.9 -e 0.001 --kmer-per-seq 100
~~~

Standard clustering:

~~~
MMseqs2 cluster module, parameters: -s 4 --min-seq-id 0.3 --cov-mode 0 -c 0.7 -e
0.001 --cluster-mode 0 --cluster-reassign
~~~

Deep clustering (profiles from standard clustering step):

~~~
MMseqs2 search module (profile-sequence search): --add-self-matches -a -s 4 -e
0.001 --cov-mode 0 -c 0.60
MMseqs2 clust module, parameters: --cluster-mode 0
~~~

Multiple sequence alignments were generated using MUSCLE, or if the cluster was too big or the alignment too large, MMseqs2 alignment against central sequence of the cluster was used.

HHmake with option -M50 was used to generate the HMMs.

### SI2: Pseudo-count computation

The viral proportion Vp can suffer from little representation of viral proteins in databases. For instance if a protein P is identified in a single viral clade C, and C contains only one viral genome in the reference database, then a naïve computation of Vp as the ratio of number of genomes in the clade containing the protein P over the total number of genomes in the clade would lead to a Vp of 1.

Adding prior information in the context of small data is crucial to have a reliable measure of the viral proportion. A standard way to cope with this issue is simply to add pseudo counts to the number of genomes in the clade. These pseudo counts vanish relatively to the real count as soon as the reference database contains enough viral genomes in the clade.

We decided to calibrate the pseudo counts so that one ends up with a reliable assessment of the “viralness” in the extreme case of a single representative genome in the clade. In particular, we want the viral quotient to be less than 0.8 as soon as the hostness (=Cp) is greater or equal to 1%, which would make the category of the protein either Cellular, Shared or Unknown according to the classification scheme described in the subsection “Material & Methods - Estimating the viralness of each enVhog” of the main paper.

Let be the pseudocount parameter. For a protein of interest P, when there is a single homolog in the viral reference database, with a single viral genome in its clade, one has:

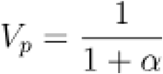

In this case, as soon as the cellular proportion of the protein is non-trivial (say Cp >1%), we want the VQ to be low enough not to classify the protein as viral. According to our scheme, VQ has to be lower than 0.8. This induces a lower bound constraint on the pseudocount.

Indeed, if the hostness is > 1%, the viral quotient is:

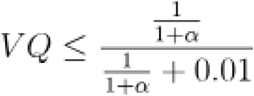

If we want to upper bound the viral quotient by 0.8, then we have the following equation:

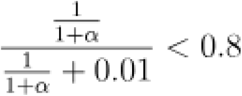

Which is equivalent to:

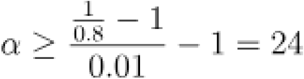

We therefore set the pseudocount parameter to 24.

When homologs of the protein of interest are present in different clades, the pseudocount parameter is equally spread among the clades, hence ending up with the formula:

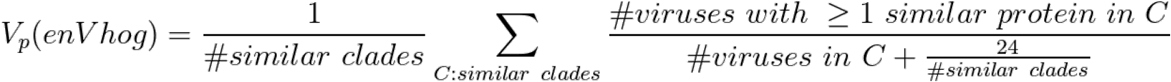

### SI3: Iterative class assignment

#### 1 Assignation of categories to contigs

As most of the proteins have been classified as Unknown in the previous section, we leveraged co-localization information of clusters on contigs to tag contigs as either Viral, Cellular (contamination), Hybrid or Unknown.

We assigned each contig to a class according to the following (stringent) criteria:

1. If the contig contains a majority of protein of category V (i.e. #V > #C+#S+#U on this contig) -> tag the contig as Viral
2. If the contig contains a majority of protein of category C (i.e. #C > #V+#S+#U on this contig) -> tag the contig as Cellular
3. If the contig contains a majority of protein of category U (i.e. #U > #V+#S+#C on this contig) -> tag the contig as Unknown
4. Else it is tagged as H (hybrid contig, containing a mixture V, C and U proteins)

Note that for contigs there is no shared (S) class as they are either from a viral or bacterial genome.

Then we can leverage the contig classes to annotate the proteins on the newly labeled contigs.

#### 2 Bayesian modeling of the contig counts

For a given enVhog, let N be the number of annotated contigs (of category V/C/H) carrying this enVhog. We model the number of contigs n_V_, n_C_, n_H_ (N=n_V_ + n_C_ + n_H_) broke down by category as a multinomial distribution parameterized by p_V_, p_C_, p_H_:

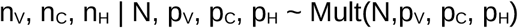

More precisely, p_V_, p_C_, p_H_ depend only on the category (V/S/C/U) of the enVhog. So if k is the category of the considered enVhog, one gets:

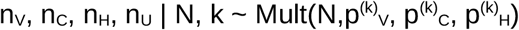

Rigorously, by setting an uninformative Dirichlet prior distribution on 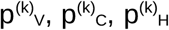 (for each k in {V,S,C,U}), one gets a Dirichlet posterior distribution on 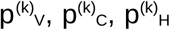 when seeing the data for all enVhogs of each category. Since there is a large number of enVhogs, the posterior distribution is sharply peaked around its mode, and the posterior distribution of the 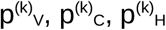 is constant to:

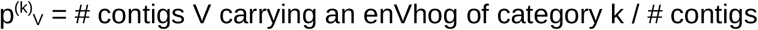

And similarly for other contig categories C and H.

Thus, after having inferred this way the 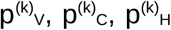 for each category k in {V,S,C,U}, we are able to compute the likelihood for a given enVhog to be in the category k:

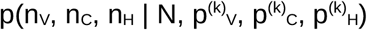

In particular, we can compute the ratio:

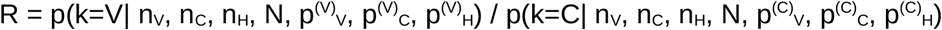

Using Bayes’ rule:

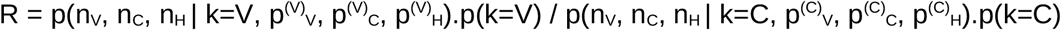

By replacing with the multinomial distribution formula, we simply get:

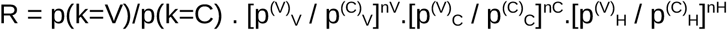

Where the prior p(k=V) is straightforwardly inferred from the total counts:

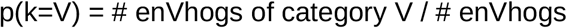

And similarly for p(K=C).

## 3 Iterative algorithm for assigning enVhog categories

For each uncertain enVhog (i.e. of category S or U), we computed the R ratio of V and C categories and apply the following decision:

1. If R>5, the enVhog is labeled as V
2. If R<1/5, the enVhog is labeled as C
3. Else the category remained unchanged (S or U) since we do not have enough evidence to decide on the category

Finally, for enVhogs labeled as C, we force them back to category S if their hostness is below 5% (indeed indicating that the protein is not well conserved in cellular genomes).

After the new category has been set to each envhog, we recomputed the contig categories using the scheme described in the previous section a.

We ran 3 iterations of this scheme, recomputing a fresh ratio R based on the contig and enVhog categories of the previous iteration.

## Notes

http://envhog.u-ga.fr/envhog/

### Competing Interest Statement

The authors have declared no competing interest.

### Summary of Updates

The manuscript has been recommended by PCI Micriobiology, and final editing has been done to comply with PCJ style.

